# Giant *GAL* gene clusters for the melibiose-galactose pathway in *Torulaspora*

**DOI:** 10.1101/2020.09.09.289694

**Authors:** Anjan Venkatesh, Anthony L. Murray, Aisling Y. Coughlan, Kenneth H. Wolfe

**Affiliations:** UCD Conway Institute and School of Medicine, University College Dublin, Dublin 4, Ireland

## Abstract

In many yeast species the three genes at the center of the galactose catabolism pathway, *GAL1*, *GAL10* and *GAL7*, are neighbors in the genome and form a metabolic gene cluster. We report here that some yeast strains in the genus *Torulaspora* have much larger *GAL* clusters that include genes for melibiase (*MEL1*), galactose permease (*GAL2*), glucose transporter (*HGT1*), phosphoglucomutase (*PGM1*), and the transcription factor *GAL4*, in addition to *GAL1*, *GAL10*, and *GAL7*. Together, these 8 genes encode almost all the steps in the pathway for catabolism of extracellular melibiose (a disaccharide of galactose and glucose). We show that a progenitor 5-gene cluster containing *GAL 7-1-10-4-2* was likely present in the common ancestor of *Torulaspora* and *Zygotorulaspora*. It added *PGM1* and *MEL1* in the ancestor of most *Torulaspora* species. It underwent further expansion in the *T. pretoriensis* clade, involving the fusion of three progenitor clusters in tandem and the gain of *HGT1*. These giant *GAL* clusters are highly polymorphic in structure, and subject to horizontal transfers, pseudogenization and gene losses. We identify recent horizontal transfers of complete *GAL* clusters from *T. franciscae* into one strain of *T. delbrueckii*, and from a relative of *T. maleeae* into one strain of *T. globosa*. The variability and dynamic evolution of *GAL* clusters in *Torulaspora* indicates that there is strong natural selection on the *GAL* pathway in this genus.

## Introduction

Physical clusters of genes that function in the same process or metabolic pathway are relatively rare in yeasts (Riley et al., 2016; Rokas et al., 2018), but in budding yeasts (Saccharomycotina) the known examples include gene clusters for the pathways *NIT* (nitrate assimilation (Ávila et al., 2002)), *PUL* (pulcherrimin synthesis (Krause et al., 2018)), *NAG* (N-acetyl glucosamine catabolism (Yamada-Okabe et al., 2001)), *LAC* (lactose utilization (Varela et al., 2019)), *DAL* (allantoin degradation (Wong and Wolfe, 2005)), *MAL* (maltose utilization (Viigand et al., 2018)), and *GAL* (galactose utilization (Slot and Rokas, 2010)). The *GAL* pathway is one of the most intensively studied systems in yeast genetics. The canonical *GAL* gene cluster was first characterized in *Saccharomyces cerevisiae*, where it consists of three genes (*GAL1*, *GAL10* and *GAL7*) that code for the pathway to convert intracellular β-D-galactose to glucose-1-phosphate (Fig. 1) (Douglas and Hawthorne, 1964; St John and Davis, 1981). The same three genes are clustered in the same order in *Kluyveromyces lactis* (Webster and Dickson, 1988) and most other species in the family Saccharomycetaceae. A similar cluster of *GAL 1-10-7*, interspersed with two genes of unknown function, occurs in *Candida albicans* and other species in the CUG-Ser1 clade (Slot and Rokas, 2010). In more divergent yeasts the *GAL* genes are generally not clustered, except for four genera (*Schizosaccharomyces*, *Nadsonia*, *Brettanomyces* and *Wickerhamomyces*) that gained clusters by horizontal transfer from donors in the CUG-Ser1 clade, and two genera (*Cryptocococcus* and *Lipomyces*) in which *GAL* clusters appear to have formed independently (Slot and Rokas, 2010; Haase et al., 2020).

**Figure 1.**
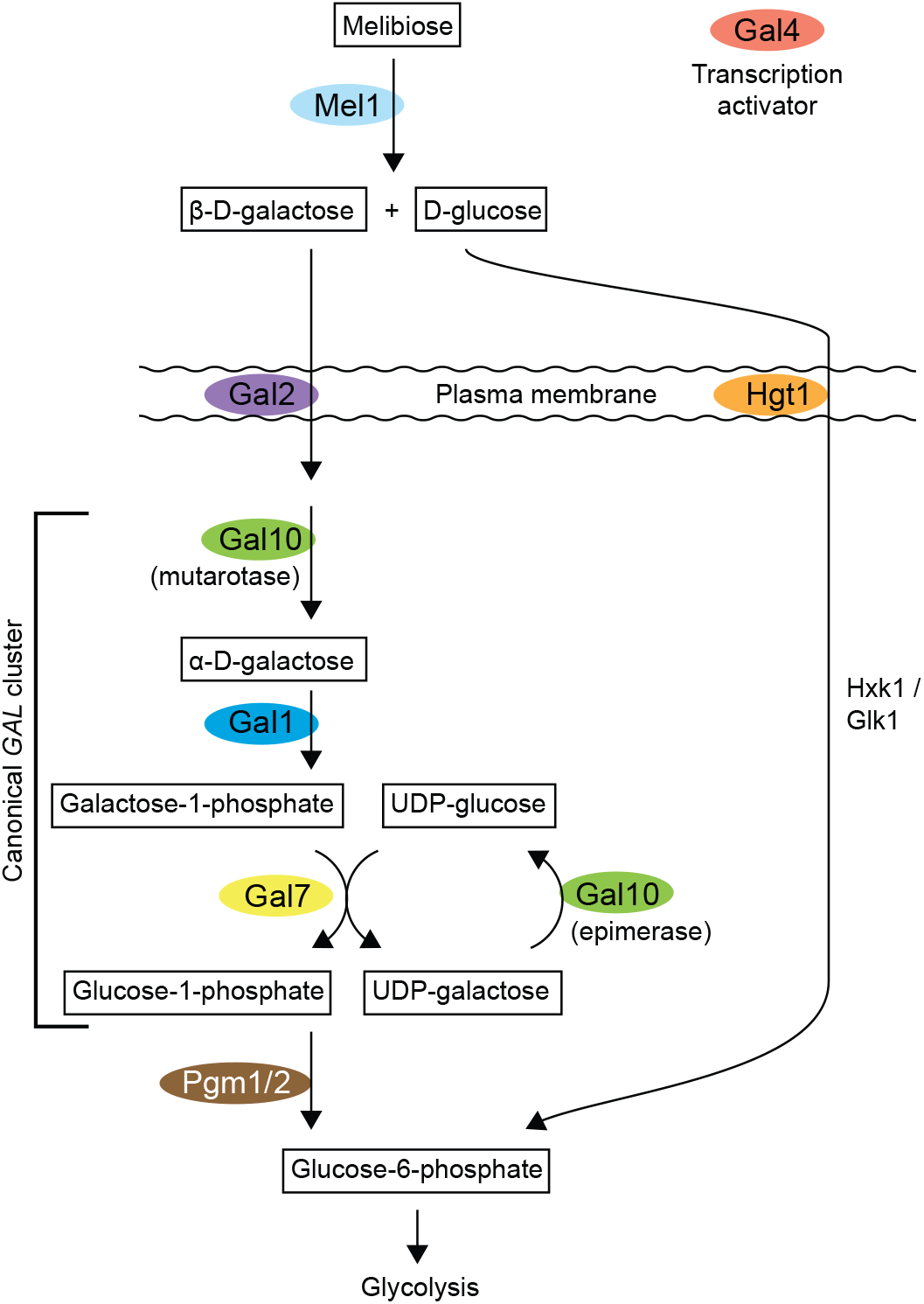
The yeast biochemical pathway for catabolism of extracellular melibiose (Holden et al., 2003). Colored backgrounds indicate genes that are located in clusters in *Torulaspora* species. Gal10 has two distinct functions, mutarotase and epimerase, performed by two domains of the protein. Hgt1 has been reported to transport galactose as well as glucose in *K. lactis* (Baruffini et al., 2006).

It is widely thought that clustering of metabolic genes evolves as a mechanism for co-regulating the expression of genes, and that clustering can be selected for if an intermediate metabolite in the pathway is toxic – as is the case for galactose-1-phosphate in the *GAL* pathway – so that is it important to coordinate synthesis and removal of the toxin (McGary et al., 2013). The local order of genes within clusters often varies among species (Wong and Wolfe, 2005; Slot and Rokas, 2010; Naseeb and Delneri, 2012), and it is common to find that genes that are in a cluster in one species are completely absent from the genome in others (Hittinger et al., 2004; Wolfe et al., 2015). It is also common to find that the metabolic pathways encoded by clustered genes show presence/absence polymorphism within a species: for example, the *GAL* genes (including the *GAL 1-10-7* cluster but also the unclustered genes *GAL4*, *GAL2* and *GAL80*) are intact in some populations of *S. kudriavzevii* but pseudogenes in others (Hittinger et al., 2010).

We previously reported that the genome sequence of the type strain of *Torulaspora delbrueckii* (CBS1146^T^) contains a large cluster of *GAL* genes, occupying 22 kb near a telomere of chromosome 5 (Wolfe et al., 2015). As well as *GAL10* (2 copies), *GAL1* (2 copies) and *GAL7* (1 copy), the cluster also contained predicted genes *MEL1* (melibiase), *GAL2* (galactose permease), *PGM1* (phosphoglucomutase), *GAL4* (transcription factor) and *HGT1* (high-affinity glucose transporter, orthologous to *K. lactis HGT1* (Billard et al., 1996)). The genes in this cluster appeared to code for additional steps in the *GAL* pathway, both upstream and downstream of the steps encoded by the canonical *GAL1-10-7* cluster (Fig. 1). In the extended pathway, extracellular melibiose (a disaccharide) is hydrolyzed into its constituent monosaccharides β-D-galactose and D-glucose by secreted Mel1 enzyme (melibiase, an α(1,6)-galactosidase). The monosaccharides are then imported across the plasma membrane by Gal2 (for galactose) and Hgt1 (for glucose). The galactose is processed by the Gal10, Gal1 and Gal7 enzymes to yield glucose-1-phosphate, which is then converted to glucose-6-phosphate by Pgm1. A second molecule of glucose-6-phosphate is made by importing the glucose and phosphorylating it by hexokinase (Hxk1) or glucokinase (Glk1). The two molecules of glucose-6-phosphate then enter the glycolytic pathway. Thus, the *T. delbrueckii* gene cluster appeared to contain genes for all the steps needed to convert melibiose into two molecules of glucose-6-phosphate, except for hexokinase/glucokinase; there are *HXK1* and *GLK1* genes in the *T. delbrueckii* genome but they are not in the cluster. The *T. delbrueckii* cluster also contains an ortholog of *S. cerevisiae GAL4*, the transcription factor that positively regulates expression of the other *GAL* genes (Hittinger et al., 2004).

In this study, we used genome sequences from additional species and strains of *Torulaspora*, generated in other studies (Galeote et al., 2018; Shen et al., 2018; Coughlan et al., 2020), to investigate the origin and evolution of *GAL* clusters in *Torulaspora* and related genera. We find that the large *GAL* cluster in the type strain of *T. delbrueckii* is atypical of this species, because all 14 other *T. delbrueckii* strains that we examined have no cluster, and we show that the cluster in the type strain of *T. delbrueckii* was acquired from *T. franciscae* recently by horizontal gene transfer. We also uncovered an extraordinary diversity of allelic *GAL* gene cluster structures in *T. pretoriensis*, and a rich history of cluster expansion, fusion, and degeneration.

## Results

### Phylogeny and phenotypes

We examined genome sequences from multiple strains of *T. delbrueckii*, *T. pretoriensis* and *T. globosa*, and from single strains of other *Torulaspora* species, as well as *Zygotorulaspora mrakii*, *Zygotorulaspora florentina*, *Zygosaccharomyces rouxii*, *Kluyveromyces lactis* and *S. cerevisiae*. The phylogeny of the species, and a summary of the major events we infer to have occurred during *GAL* cluster evolution in *Torulaspora*, is shown in Figure 2. One gene in the well-known *GAL* system of *S. cerevisiae*, *GAL3*, is a paralog of *GAL1* that was formed by the whole-genome duplication (WGD). *Torulaspora* and all the other genera considered here diverged from *S. cerevisiae* before the WGD occurred, so their *GAL1* genes are orthologous to both *GAL1* and *GAL3* in *S. cerevisiae*. Another gene, *GAL80*, coding for a corepressor of *GAL* gene expression, is absent from most *Torulaspora* species (Fig. 2).

**Figure 2.**
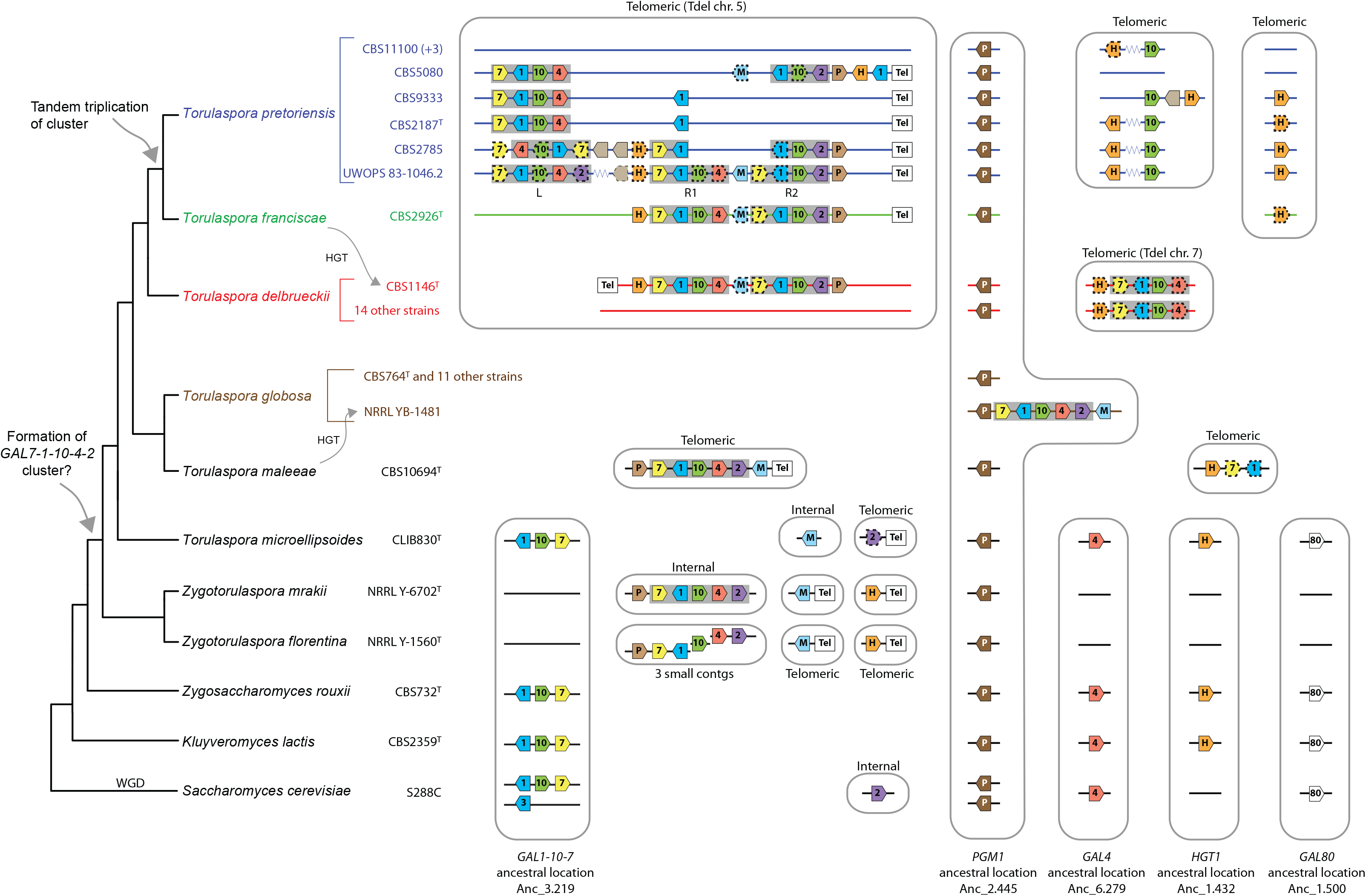
Synteny relationships among *GAL* genes and clusters in *Torulaspora* species and outgroups. Genes are labeled with their *GAL* gene number (7, 1, 10, 4, 2, or 80), or M (*MEL1*), P (*PGM1*), or H (*HGT1*). Dashed borders on gene symbols indicate pseudogenes. Gray backgrounds highlight groups of adjacent genes with the progenitor cluster gene order *GAL 7-1-10-4-2* or subsets thereof. Large gray boxes indicate groups of genes that are at syntenic locations in different strains/species, and are indicated as being either telomeric or internal to chromosomes. Ancestral gene locations refer to the numbering system of Gordon et al. (2009) and are internal to chromosomes. Different P symbols are used to distinguish between *PGM1* genes at the ancestral location (*PGM1_anc*, dark brown), and duplicate *PGM1* genes in *GAL* clusters (*PGM1_dup*, light brown). Tel indicates a region inferred to be close to a telomere (subtelomeric), and zigzag symbols in *T. pretoriensis* indicate intervening regions of 10-15 kb with no genes related to *GAL* metabolism. The tree topology is from the phylogenomic analysis of Shen et al. (2018) with *T. globosa* added as in (Saluja et al., 2012; Kaewwichian et al., 2020).

A *GAL* cluster is present in at least some strains of all the *Torulaspora* species we studied. We tested the ability of several strains to grow on solid media containing galactose, melibiose, or glucose as a sole carbon source (Fig. 3). We found that the ability to grow on galactose correlates with the presence of intact copies of the genes *GAL1*, *GAL10* and *GAL7* in the genome, and the ability to grow on melibiose correlates with the presence of an intact *MEL1* gene (Fig. 3). The starting point for our study was the large *GAL* cluster on chromosome 5 of *T. delbrueckii* strain CBS1146^T^ (Wolfe et al., 2015), and we found that this strain can grow on galactose whereas *T. delbrueckii* strain L09, which lacks the cluster, cannot (Fig. 3). However, we were surprised to find that *T. delbrueckii* CBS1146^T^ cannot grow on melibiose despite apparently having a *MEL1* gene. We realized that the open reading frame we originally annotated as *MEL1* (*TDEL0E00170*) is truncated at the 5’ end relative to other *MEL1* genes. Comparison to a functional *MEL1* gene previously characterized by Oda and Fukunaga (1999) from *T. delbrueckii* strain IFO1255 shows that CBS1146^T^ has a TGG (Trp) -> TGA (stop) mutation at codon 38 which removes the region coding for the secretion signal, so the *MEL1* gene of CBS1146^T^ is a pseudogene. A second discrepancy between genotypes and phenotypes occurs in *T. pretoriensis* CBS2187^T^, which grows poorly on galactose despite containing *GAL1*, *GAL10* and *GAL7* genes (Fig. 3). This discrepancy is discussed later.

**Figure 3.**
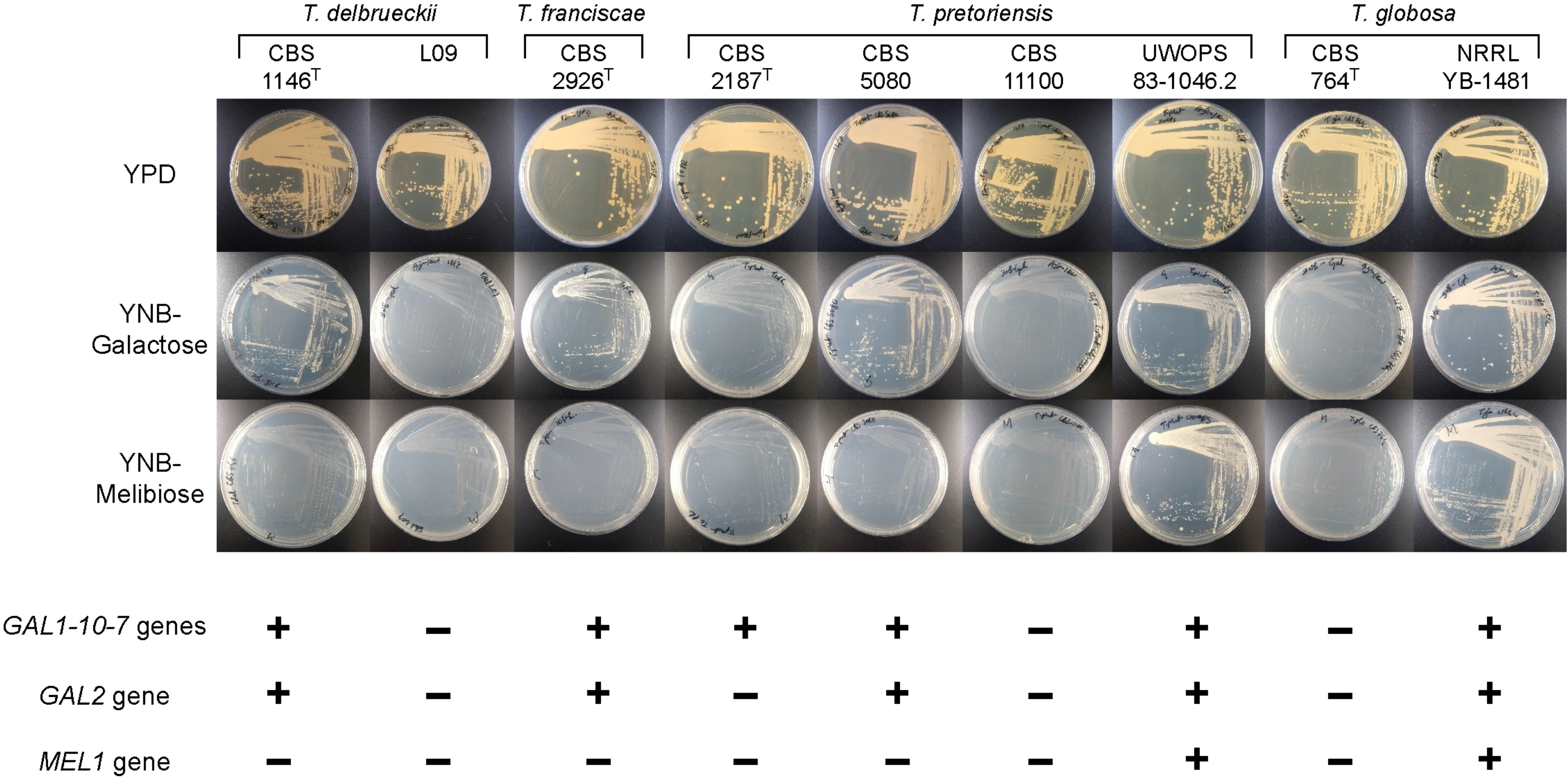
Growth of *Torulaspora* strains on galactose, melibiose, and glucose (YPD) media. Plates were incubated at 30° C for 48 hours before photographing. The lower panel indicates the presence or absence of intact genes in each genome.

### Synteny relationships

Synteny comparisons among the *Torulaspora* species and outgroups revealed a complex pattern of relationships and gene relocations (Fig. 2). For some loci, we refer to the Ancestral gene numbering system of Gordon et al. (2009), which numbers genes sequentially along the 8 chromosomes inferred to have existed just prior to the WGD, for example locus Anc_8.123 is the 123^rd^ gene along Ancestral chromosome 8. This numbering system is also used in our Yeast Gene Order Browser (ygob.ucd.ie) (Byrne and Wolfe, 2005).

In the outgroup species shown at the bottom of Figure 2 (*S. cerevisiae*, *K. lactis*, *Z. rouxii*), the only genes in the *GAL* pathway that are clustered are *GAL1*, *GAL10* and *GAL7*, and they occur in the order *GAL 1-10-7*. This arrangement is conserved in *T. microellipsoides*, including the flanking genes *SNQ2* and *RPT2* (Anc_3.216 to Anc_3.220). This cluster is at an internal chromosomal site in these species, i.e. it is not subtelomeric. In the outgroups, the other genes in the pathway are at conserved, dispersed, places in the genome (*PGM1* = Anc_2.445; *GAL4* = Anc_6.279; *HGT1* = Anc_1.432; *GAL80* = Anc_1.500), and *MEL1* is not present at all.

### Formation of a large *GAL* cluster in the common ancestor of *Torulaspora* and *Zygotorulaspora*

In *Zygotorulaspora mrakii*, the cluster has expanded to 6 genes: it contains *GAL 7-1-10-4-2* and a *PGM1* gene (Fig. 2). *Z. mrakii* also has an unlinked *MEL1* gene, which was previously shown to be functional by Oda and Fujisawa (2000). The 6-gene cluster has gained genes for the pathway steps upstream (*GAL2*) and downstream (*PGM1*) of the steps encoded by the 3-gene cluster, as well as gaining the transcription factor *GAL4*. It is interesting that the order of the 3 genes has also changed, from *GAL 1-10-7* in the outgroups to *GAL 7-1-10* in *Z. mrakii*. The *Z. mrakii* 6-gene cluster is located at an internal chromosomal site between *EST3* (Anc_7.128) and *URM1* (Anc_7.129). The cluster therefore appears to have become inserted between two genes that were ancestrally neighbors. In the genome assembly of a second *Zygotorulaspora* species, *Z. florentina* (accession number PPJY02000000), the same six genes are found on three small contigs: one containing only *PGM1-GAL7-GAL1*, one containing only *GAL10*, and one containing only *GAL4-GAL2*, so it is unclear whether *Z. florentina* has a *GAL* cluster organization identical to that in *Z. mrakii* or a more fragmented organization.

In *T. maleeae*, there is a 7-gene cluster with identical gene order to the 6-gene cluster of *Z. mrakii*, plus *MEL1* (Fig. 2). This cluster appears to be at a subtelomeric location, and the *EST3* and *URM1* genes (Anc_7.128/7.129) are adjacent in *T. maleeae*. Both *T. maleeae* and the two *Zygotorulaspora* species have two *PGM1* genes. The first, designated *PGM1_anc*, is at the ancestral *PGM1* location (Anc_2.445). It is syntenic with the *PGM1* genes of other yeasts, including the *PGM1/PGM2* gene pair of *S. cerevisiae*, which is a WGD pair. The second, designated *PGM1_dup*, is a duplicated copy of *PGM1* located in the *GAL* cluster.

The gene order *GAL 7-1-10-4-2*, as seen in *Z. mrakii* and *T. maleeae*, is a pattern that recurs throughout the *GAL* clusters of most *Torulaspora* species that will be described in the following sections. However, *T. microellipsoides* has an ancestral-type cluster (*GAL 1-10-7*) at the ancestral location (Anc_3.219), rather than the *GAL 7-1-10-4-2* pattern, even though phylogenomic analysis (Shen et al., 2018) has indicated that the genus *Torulaspora* is monophyletic and *Zygotorulaspora* is an outgroup to it. *T. microellipsoides* also has a *MEL1* gene at an unlinked, non-telomeric location (Fig. 2).

The organization of *GAL* genes in *T. microellipsoides* resembles the outgroup species more closely than it resembles other *Torulaspora* species, whereas the *Z. mrakii* organization resembles *Torulaspora* species (Fig. 2). In phylogenetic trees of individual *GAL* genes, *T. microellipsoides* is often placed outside *Zygotorulaspora* (Fig. 4), in contrast to the phylogenomic tree. Moreover, *GAL80* is present in *T. microellipsoides* but absent in the other *Torulaspora* species and in *Zygotorulaspora* (Fig. 2). Together, these results suggest that the phylogenomic tree might be incorrect regarding the branching order of *Zygotorulaspora* and *T. microellipsoides*. Alternatively, there may have been horizontal transfer of a *GAL* cluster between the *Zygotorulaspora* and *Torulaspora* branches in either direction, after *T. microellipsoides* diverged from the rest of the genus *Torulaspora*, making the *GAL* phylogeny different from the phylogeny of the rest of the genome.

**Figure 4.**
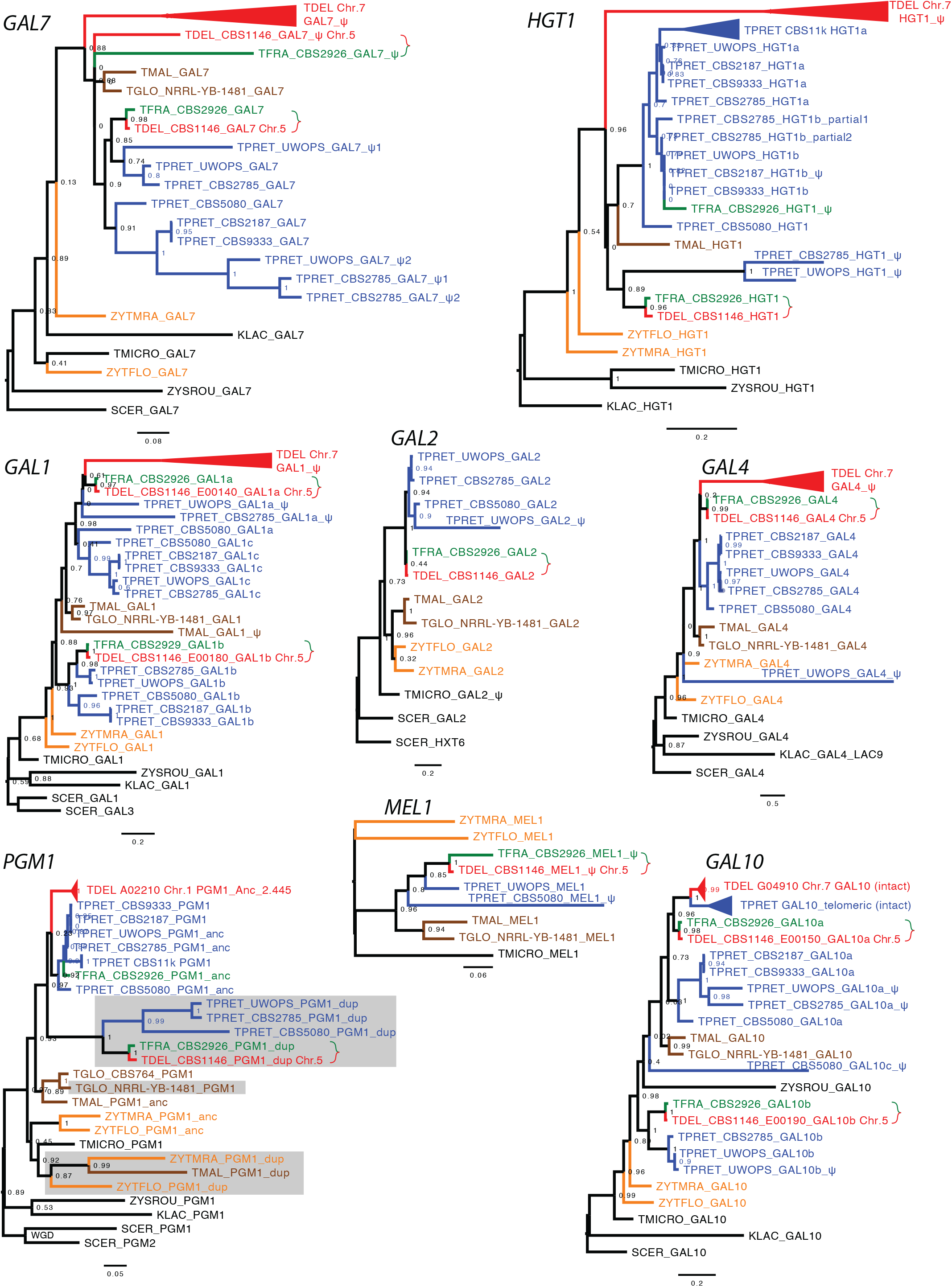
Phylogenetic trees of *GAL*, *PGM1*, *HGT1* and *MEL1* genes. Branches are colored by species. Some groups of closely related sequences have been collapsed (triangles). Green/red braces mark gene pairs showing horizontal transfer between *T. franciscae* (TFRA) and *T. delbrueckii* (TDEL) strain CBS1146^T^. In the *PGM1* tree, gray rectangles indicate genes that are located in *GAL* clusters, and for genomes with two *PGM1* genes the copies are labeled *PGM1_anc* and *PGM1_dup*; other genomes have only one gene. Approximate likelihood ratio test (aLRT) branch support values are shown.

In summary, the point of origin of the *GAL 7-1-10-4-2* cluster pattern is not fully clear, but it appears to have been present in the common ancestor of the genera *Zygotorulaspora* and *Torulaspora*. It is first seen with *PGM1* at one end, and later gained *MEL1* at the other end.

### Horizontal *GAL* cluster transfer into one strain of *T. globosa*

*T. globosa* is a sister species to *T. maleeae*. We sequenced the genomes of 12 strains of *T. globosa* (Coughlan et al., 2020 and A.Y.C. and K.H.W., unpublished) and found that 11 of them, including the type strain CBS764^T^, have no *GAL* genes. However, one strain, *T. globosa* NRRL YB-1481, has a *GAL* cluster, and the organization of this cluster is very similar to the *T. maleeae* cluster (Fig. 2). Phylogenetic trees of *GAL 7*, *1*, *10*, *4*, *2* and *MEL1* all show that the *T. globosa* NRRL YB-1481 genes group with the *T. maleeae* genes (Fig. 4). In plate tests, *T. globosa* NRRL YB-1481 was able to grow on melibiose and galactose, whereas *T. globosa* CBS764^T^ could not (Fig. 3).

Interestingly, the *GAL* cluster in *T. globosa* strain NRRL YB-1481 has formed at the ancestral location of *PGM1* (Anc_2.445; Fig. 2). This strain has only one *PGM1* gene, in contrast to *T. maleeae* and *Z. mrakii* which have two (*PGM1_anc* and *PGM1_dup*). Since most *T. globosa* strains have no *GAL* genes, the most plausible scenario to explain the presence of a cluster in NRRL YB-1481 is that it originated by horizontal transfer. In view of the relatively low DNA sequence identity (74%) between the *T. globosa* NRRL YB-1481 and *T. maleeae* clusters, the donor is more likely to have been an unidentified species related to *T. maleeae*/*T. globosa*, rather than *T. maleeae* itself.

Although it is possible that recombination between the *PGM1* genes in the donor cluster and the recipient *T. globosa* NRRL YB-1481 genome might have guided integration of the cluster, this seems unlikely because the *T. maleeae* and *T. globosa PGM1* genes are currently in opposite orientations relative to their neighbor *GAL7* (Fig. 2). Also, a phylogenetic tree of *PGM1* sequences (Fig. 4) places the single, cluster-associated, *PGM1* of *T. globosa* NRRL YB-1481 at the position expected for a *PGM1_anc* gene: it is in a clade with the single *PGM1* gene of *T. globosa* CBS764T and *T. maleeae PGM1_anc*, and far away from *T. maleeae PGM1_dup* which lies in a clade with *PGM1_dup* genes from *Z. mrakii* and *Z. florentina*.

### Horizontal *GAL* cluster transfer from *T. franciscae* into *T. delbrueckii*

T. pretoriensis, *T. franciscae* and *T. delbrueckii* form a clade of three species whose *GAL* clusters, when present, are greatly expanded and contain numerous *GAL* pseudogenes as well as functional genes. We analyzed data from multiple strains of *T. delbrueckii* and *T. pretoriensis*, but we have only one genome sequence from *T. franciscae* (the type strain, CBS2926^T^).

In the set of 15 *T. delbrueckii* strains that we analyzed, none except CBS1146^T^ contains a *GAL* cluster, which suggests that the cluster was gained by horizontal transfer. The CBS1146^T^ cluster is identical in gene organization to a cluster in the type strain of *T. franciscae*, and the two clusters have 97% DNA sequence identity over 22 kb. The similarity between these two species is much higher than between either of them and *T. pretoriensis*, even though *T. pretoriensis* is a sister species to *T. franciscae* (Fig. 2). Phylogenetic trees from individual genes in the cluster consistently place *T. delbrueckii* CBS1146^T^ beside *T. franciscae* (Fig. 4). We therefore infer that horizontal transfer occurred from *T. franciscae* to *T. delbrueckii*. Curiously, although the cluster is near a telomere in both species, the two species have opposite orientations of the cluster relative to the telomere (Fig. 2).

The *MEL1* genes in the clusters in the type strains of both *T. franciscae* and *T. delbrueckii* are pseudogenes, and these strains are unable to grow on melibiose but able to grow on galactose (Fig. 3). In a previous study by Oda and Tonomura (1996), 12 of 28 *T. delbrueckii* strains examined, including the type strain, were found to be able to grow on galactose. Only one of the *T. delbrueckii* strains (IFO 1255) could grow on melibiose as well as galactose and was shown to have an intact *MEL1* gene (Oda and Tonomura, 1996; Oda and Fukunaga, 1999).

### Extensive structural polymorphism of *T. pretoriensis GAL* clusters

We analyzed genome sequences from nine strains of *T. pretoriensis*, of which five have large and variable *GAL* clusters, and the other four have none. The four strains without clusters (CBS11100, CBS11121, CBS11123, CBS11124) are closely related to each other, so only CBS11100 is shown in Figure 2. Among the five strains with clusters, there is extensive structural polymorphism, with only two strains (CBS2187^T^ and CBS9333) having similar organization. All the *GAL* clusters in *T. pretoriensis* strains appear to be near telomeres.

The most complex *GAL* cluster in *T. pretoriensis* is in strain UWOPS 83-1046.2 (Fig. 2; we refer to this strain hereafter as UWOPS). It spans 42 kb and contains 8 intact genes and 8 pseudogenes related to galactose metabolism. It also contains 2 unrelated genes and 1 unrelated pseudogene, which appear to be of subtelomeric origin. These unrelated genes occupy a region of 15 kb inside the cluster and divide it into two parts, left and right. The right part is almost identical in gene organization to the large *GAL* cluster that was transferred between *T. franciscae* and *T. delbrueckii* CBS1146^T^, the only differences being some genes that are pseudogenes in *T. pretoriensis* UWOPS but intact in *T. franciscae* and *T. delbrueckii* CBS1146^T^, or vice versa (*HGT1*, *MEL1*, and one copy each of *GAL1* and *GAL10*; Fig. 2). Phylogenetic analysis of the genes in this region (Fig. 4) shows that, in all cases, *T. franciscae* and *T. delbrueckii* CBS1146^T^ form a clade with *T. pretoriensis* UWOPS outside, which contradicts the expected species phylogeny (Fig. 2) and supports the hypothesis of horizontal transfer between *T. franciscae* and *T. delbruckii*.

We tested the phenotypes of four *T. pretoriensis* strains (Fig. 3). As expected, only UWOPS can grow on melibiose – it is the only strain with intact *MEL1*. On galactose, CBS11100 cannot grow (it has no *GAL* cluster), CBS5080 and UWOPS grow well, and the type strain CBS2187^T^ grows more slowly. The poor growth of the type strain of *T. pretoriensis* on galactose is consistent with previous studies. Oda and colleagues reported that fermentation of galactose or melibiose by strain YK-1, which is a non-sedimenting derivative of *T. pretoriensis* CBS2187T (syn. IFO 10218), was undetectable after 2 days, whereas *T. pretoriensis* CBS5080 (IFO 0022) and *T. franciscae* CBS2926^T^ (IFO 1360) fermented galactose but not melibiose (Oda and Tonomura, 1993; Oda and Tonomura, 1996). Oda’s results are consistent with our results in Figure 3, except that we find that growth of CBS2187^T^ on galactose is slow rather than absent. A possible reason for the poor growth is that there is no *GAL2* galactose transporter gene anywhere in the *T. pretoriensis* CBS2187^T^ genome; it is the only strain tested in Figure 3 that has the *GAL* enzyme genes without the transporter gene.

### Cluster expansion by tandem triplication of progenitor *GAL 7-1-10-4-2* clusters

Closer examination of the *T. pretoriensis GAL* clusters shows that they have an internal structure that is based on tandem triplication of the *GAL 7-1-10-4-2* pattern mentioned earlier. This structure is most clearly seen in *T. pretoriensis* UWOPS which has three copies of the pattern: including pseudogenes, it has *GAL 7-1-10-4-2* in the left part of the cluster, and *GAL 7-1-10-4* (without *GAL2*) followed by *GAL 7-1-10-2* (without *GAL4*) in the right part. The other genes in the cluster (*HGT1*, *MEL1*, *PGM1*, and the unrelated genes between the left and right parts) are located at the junctions between these three copies of the pattern.

This arrangement suggests that the large UWOPS cluster was formed by tandem fusion of three smaller progenitor clusters that we designate L, R1 and R2, corresponding to the left part and two sections of the right part of the current cluster (Fig. 2). We postulate that L contained *GAL 7-1-10-4-2*, R1 originally contained *HGT1 – GAL 7-1-10-4-2*, and R2 originally contained *MEL1 – GAL 7-1-10-4-2 – PGM1*. Subsequently, many of the triplicated *GAL* gene copies became pseudogenes or relics (very short pseudogenes), and no trace remains of *GAL2* in R1 or *GAL4* in R2. Notably, although there are many pseudogenes in the *T. pretoriensis* clusters (of all strains), there are no pseudogenes that indicate that *HGT1*, *MEL1*, or *PGM1* was ever duplicated within the clusters; all the duplications are of *GAL* genes. Therefore we suggest that the triple-size cluster did not arise by triplicating a single progenitor cluster, but instead arose by fusion of three progenitor clusters that were similar (containing *GAL 7-1-10-4-2*) but already different regarding their content of *HGT1*, *MEL1* and *PGM1*.

The clusters in the other *T. pretoriensis* strains are smaller than in UWOPS but still consistent with the hypothesis of cluster expansion by tandem fusion of progenitors. Strain CBS2785 has an overall organization similar to UWOPS, but it has lost *MEL1* and adjacent parts of R1 and R2. It has also sustained an inversion of *GAL1-10-4* in the L part, probably in conjunction with the formation of an extra relic of *GAL7* that is also in inverted orientation. Strain CBS5080 has parts L and R2 but not R1, and it also has additional *HGT1* and *GAL1* genes to the right of R2. Strains CBS2187^T^ and CBS9333 have only part L and an additional *GAL1* gene; they lack *MEL1*, *HGT1* and *PGM1* in the cluster and have only one *PGM1* gene in their genomes (at the ancestral locus Anc_2.445). The phylogenies of most genes and pseudogenes in the *T. pretoriensis* clusters (Fig. 4) generally support the relationships shown in Figure 2, which are based on synteny as well as phylogenetic considerations. It is impossible to infer the complete history of the *T. pretoriensis* clusters, but we can conclude that (i) at least three progenitor clusters fused in tandem to form them, and (ii) they are undergoing extensive within-species structural rearrangement and turnover.

### Vestigial *GAL* clusters and extra unclustered *GAL10* and *HGT1* genes

The large *GAL* cluster in *T. delbrueckii* originated by horizontal transfer from *T. franciscae*. Among our sequenced strains, it is only present in CBS1146^T^ and is located near a telomere of chromosome 5. However, in addition, all 15 *T. delbrueckii* strains (including CBS1146^T^) also contain an intact *GAL10* gene near a telomere of chromosome 7 (Fig. 2). It is located beside four pseudogenes in the arrangement *HGT1* – *GAL 7-1-10-4*, where *GAL10* is the only intact gene, so it appears to be a remnant of a primordial *GAL* cluster that has almost disappeared. Its structure is the same as the R1 primordial cluster inferred in *T. pretoriensis*.

Similarly, most strains of *T. pretoriensis* have an extra copy of *GAL10*, located near *HGT1* and a telomere (Fig. 2). This *GAL10* gene is present even in strains such as CBS11100 that cannot utilize galactose. Therefore, many strains of both *T. delbrueckii* and *T. pretoriensis* contain *GAL10* but no other *GAL* genes. This situation has also been seen in other yeasts (Haase et al., 2020) but its physiological significance is unknown.

An extra vestigial telomeric *GAL* cluster is also seen in *T. maleeae*, containing an intact *HGT1* gene and pseudogenes of *GAL7* and *GAL1* (Fig. 2). Thus, in both *T. maleeae* and *T. pretoriensis*, high-affinity glucose transporter function is provided by an *HGT1* gene that is neither located at the ancestral *HGT1* locus (Anc_1.432), nor in an active *GAL* cluster containing intact *GAL1* and *GAL7*, but in a remnant of a degraded cluster at a telomeric location that sometimes also includes *GAL10*. Notably, in the only *T. pretoriensis* strain that includes an intact *HGT1* in its *GAL* cluster (CBS5080), there are no additional telomeric *HGT1* or *GAL10* genes (Fig 2).

## Discussion

The *GAL* clusters of *Torulaspora* species are remarkably large and heterogeneous. There are polymorphisms both for presence/absence of the cluster, and for gene order within the cluster. Formation of pseudogenes is common. As a result, *Torulaspora* strains and species vary in their ability to grow using galactose or melibiose as the sole carbon source. It is difficult to correlate these differences with the ecology of the yeasts, because relatively little is known about their natural environments. *T. delbrueckii* and *T. microellipsoides* are frequently isolated from high-sugar anthropic environments such as food spoilage and fermented fruit juices, whereas most isolates of *T. franciscae*, *T. pretoriensis*, *T. globosa*, and *T. maleeae* come from soil (Kurtzman, 2011). For the two strains that gained *GAL* clusters by horizontal transfer, *T. globosa* NRRL YB-1481 was isolated from soil in Ghana, and the origin of the type strain of *T. delbrueckii* CBS1146^T^ is uncertain.

The cluster first expanded from a canonical 3-gene *GAL 1-10-7* structure by adding *GAL2* and *GAL4*, around the time of the common ancestor of *Torulaspora* and *Zygotorulaspora*. The synteny relationships in Figure 2 suggest that a duplicate copy of *PGM1* was then recruited into the *GAL 7-1-10-4-2* cluster, followed later by relocation of *MEL1* and then *HGT1*. However, the phylogeny of *PGM1* sequences (Fig. 4) shows that there must have been multiple separate incorporations of *PGM1* into the cluster, because the *PGM1_dup* genes in the giant *GAL* clusters of the *T. pretoriensis/T. delbrueckii*/*T. franciscae* clade originated independently of the *PGM1_dup* genes in the smaller clusters of *T. maleeae* and *Z. mrakii/Z. florentina*. Including the integration of a *GAL* cluster beside *PGM1_anc* in *T. globosa* NRRL Y-1481, there were three separate, parallel, events of incorporation of *PGM1* into *Torulaspora GAL* clusters – pointing to strong selection to incorporate it. In two *Lachancea* species a *GAL* cluster including *GAL1, GAL7* and *GAL2* has formed beside *PGM1* at its ancestral location (Kuang et al., 2018), similar to what we observe in *T. globosa* NRRL Y-1481. *PGM1* is a bottleneck gene, coding for an enzyme that integrates metabolic flux from several pathways including glycogen synthesis, trehalose synthesis and the pentose phosphate pathway as well as the *GAL* pathway, and in the genera *Saccharomyces* and *Lachancea*, regulation of *PGM1* by *GAL4* has been gained and lost multiple times (Kuang et al., 2018). We find that in the species with two *PGM1* genes (Fig. 4), the *PGM1_dup* genes in the cluster contain multiple putative Gal4 binding sites (CGG-N_11_-CCG) in their upstream regions, whereas the *PGM1_anc* genes do not. In *T. globosa* NRRL YB-1481, *PGM1* is not duplicated but has Gal4 sites in the upstream region that it shares with *GAL7* (Fig. 2). Thus, in all the clusters in the *Torulaspora* clade, a *PGM1* gene has come under the regulation of *GAL4*.

Unexpectedly, our results indicate that duplication and fusion of whole clusters, rather than duplication of individual genes, was the major mechanism of evolution of *GAL* clusters. In *T. pretoriensis*, three primordial clusters fused to form one giant cluster and many of the genes later became pseudogenes. Tandem fusion of clusters may have provided an opportunity to experiment with shuffling the gene order, by allowing different gene copies to become pseudogenes. For example, in the *T. pretoriensis* clusters, the intact gene upstream of *GAL1* can be *GAL10, GAL2*, *GAL4*, or *MEL1* (Fig. 2). Haase et al. (2020) recently identified a similar fusion of two *GAL* clusters (one ancestral and one horizontally transferred) in *Nadsonia fulvescens*.

The *Torulaspora GAL* clusters include up to eight different functional genes, comprising the whole *MEL-GAL-PGM* pathway except for hexokinase/glucokinase (Fig. 1). Since the sugar kinases also function in the pathway for catabolism of glucose monomers imported into the cell by hexose transporters, the eight genes in the cluster constitute the complete set of genes that need to be activated in the presence of melibiose or galactose, and repressed in their absence. In *K. lactis*, *HGT1* was originally described as a high-affinity glucose transporter, but it can also transport galactose and is induced by galactose (Baruffini et al., 2006).

To build clusters with eight functional genes by random genomic rearrangements, natural selection on the *GAL* metabolic pathway must be exceptionally strong in *Torulaspora*. However, we have no explanation for why selection to form clusters is stronger in *Torulaspora* than in other budding yeast genera. It seems likely that regulatory changes, involving duplication of *PGM1*, loss of *GAL80*, and movement of *GAL4* into the cluster were central to expansion of the cluster. Previous work has shown that Gal4 became the major regulator of the *GAL* pathway relatively recently, displacing Rtg1/Rtg3 in an ancestor of the family Saccharomycetaceae (Choudhury and Whiteway, 2018; Haase et al., 2020). In the *Torulaspora/Zygotorulaspora* clade, the further step of moving the *GAL4* gene into the cluster has occurred. Relocation of *GAL4* into the cluster would have enabled the Gal4 protein to evolve in concert with its binding sites in the promoters of the nearby *GAL* genes. Moreover, in the *Torulaspora/Zygotorulaspora* species (except *T. microellipsoides*), Gal4 has lost the C-terminal region for interaction with the co-repressor Gal80 (Choudhury and Whiteway, 2018), and the *GAL80* gene is absent from their genomes (Fig. 2). In each cluster, multiple putative Gal4 binding sites are present upstream of each intact *GAL* gene (except *GAL4*) as well as *PGM1* and *HGT1*, but not *MEL1*. These regulatory changes may have made the cluster almost independent of other loci in the genome, and hence made it more amenable to transfer among species.

## Methods

Yeast strains were obtained from the Westerdijk Fungal Biodiversity Institute (CBS strains), the USDA Agricultural Research Service (NRRL strains), Lallemand Inc. (L09), and M.-A. Lachance (UWOPS 83-1046.2).

For growth tests, yeast strains were streaked onto agar plates made with YPD (2% dextrose) (Formedium, catalog CCM0110), YNB (yeast nitrogen base; Sigma-Aldrich, 51483) with 2% D-(*)-galactose (Sigma-Aldrich, G0625), or YNB with 2% D-(*)-melibiose (Sigma-Aldrich, 63630). Plates were incubated at 30° C for 48 hours before photographing.

For sequencing *T. globosa* strain NRRL YB-1481, cultures were grown under standard rich-medium conditions. DNA was harvested from stationary-phase cultures by homogenization with glass beads followed by phenol-chloroform extraction and ethanol precipitation. Purified DNA was concentrated with the Genomic DNA Clean and Concentrator-10 (Zymo Research, catalog D4010). Sequencing was done by BGI Tech Solutions (Hong Kong) using Illumina HiSeq 4000 (paired end, 2 x 150 bp reads), and assembled using SPAdes version 3.11.1 (Bankevich et al., 2012). Coverage was approximately 85x. All other genome sequences are from sources cited in Coughlan et al. (2020).

*GAL* clusters were annotated manually. In the *T. franciscae* genome assembly, the large cluster was initially split into three contigs due to high similarity between the two *GAL10* genes. Its organization was inferred by manually merging scaffold 86, scaffold 87, and contig C4393.

Genes were inferred to be located in subtelomeric regions if the gene is near the end of a chromosome-sized scaffold, or if DNA sequences neighboring the gene are repeat sequences that occur only near the ends of multiple very large scaffolds, or if several neighbors of the gene are members of gene families that are often found in subtelomeric regions (Brown et al., 2010) and do not have Ancestral gene numbers (Gordon et al., 2009).

Phylogenetic trees were constructed from MUSCLE alignments of amino acid sequences, using PhyML as implemented in version 5.0 of SeaView (Gouy et al., 2010). Approximate translations of pseudogenes were made by manual annotation.

## Acknowledgments

This work was supported by the European Research Council (789341) and Science Foundation Ireland (13/IA/1910).

